# Attention Promotes the Neural Encoding of Prediction Errors

**DOI:** 10.1101/522185

**Authors:** Cooper A. Smout, Matthew F. Tang, Marta I. Garrido, Jason B. Mattingley

## Abstract

The human brain is thought to optimise the encoding of incoming sensory information through two principal mechanisms: *prediction* uses stored information to guide the interpretation of forthcoming sensory events, and *attention* prioritizes these events according to their behavioural relevance. Despite the ubiquitous contributions of attention and prediction to various aspects of perception and cognition, it remains unknown how they interact to modulate information processing in the brain. A recent extension of predictive coding theory suggests that attention optimises the expected precision of predictions by modulating the synaptic gain of prediction error units. Since prediction errors code for the *difference* between predictions and sensory signals, this model would suggest that attention increases the selectivity for *mismatch information* in the neural response to a surprising stimulus. Alternative predictive coding models proposes that attention increases the activity of prediction (or ‘representation’) neurons, and would therefore suggest that attention and prediction synergistically modulate selectivity for *feature information* in the brain. Here we applied multivariate forward encoding techniques to neural activity recorded via electroencephalography (EEG) as human observers performed a simple visual task, to test for the effect of attention on both mismatch and feature information in the neural response to surprising stimuli. Participants attended or ignored a periodic stream of gratings, the orientations of which could be either predictable, surprising, or unpredictable. We found that surprising stimuli evoked neural responses that were encoded according to the difference between predicted and observed stimulus features, and that attention facilitated the encoding of this type of information in the brain. These findings advance our understanding of how attention and prediction modulate information processing in the brain, and support the theory that attention optimises precision expectations during hierarchical inference by increasing the gain of prediction errors.

## Introduction

Perception is believed to arise from a process of active inference [1], during which the brain retrieves information from past experiences to build predictive models of likely future occurrences and compares these predictions with incoming sensory evidence [2,3]. In support of the idea that prediction increases the efficiency of neural encoding, previous studies have demonstrated that predicted visual events typically evoke smaller neural responses than surprising events (e.g. evoked activity measured in terms of changes in electrical potential or blood oxygen level dependent (BOLD) response; for a review, see [4]). Interestingly, recent studies have shown that selective attention can increase [5] or reverse [6] the suppressive effect of prediction on neural activity, suggesting that attention and prediction facilitate perception [7] via synergistic modulation of bottom-up sensory signals [8–11]. It remains unclear, however, what *type of information* is modulated in the interaction between attention and prediction. This question is important because different predictive coding models make distinct predictions about how information is transmitted through the cortical hierarchy [3,8,12,13]. Here, we used multivariate forward encoding analyses to assess selectivity for two distinct types of information in the neural response to surprising stimuli – *feature* and *mismatch information* - and to test the effect of attention on these two informational codes.

A prominent version of predictive coding theory claims that top-down prediction signals ‘cancel out’ bottom-up sensory signals that match the predicted content, leaving only the remaining prediction error to propagate forward and update a model of the sensory environment [2,8,9]. Since error propagation is thought to be associated with superficial pyramidal cells [9], and these cells are thought to be primarily responsible for generating EEG signals [14,15], this theory predicts that surprising events will increase the selectivity of EEG responses to the *difference* between predicted and observed stimulus features, i.e. mismatch information. Furthermore, a recent extension of this theory suggests that selective attention optimises the expected precision of predictions by modulating the synaptic gain (post-synaptic responsiveness) of prediction error units [8] – that is, neurons coding for behaviourally relevant prediction errors should be more responsive than those coding for irrelevant prediction errors. On this account, attention should further increase selectivity for mismatch information in the neural response to surprising stimuli relative to unsurprising stimuli. Here we call this account the *mismatch information model*.

Alternative predictive coding models [12,13,16] propose that predictions – as opposed to prediction errors – are propagated forward through the visual hierarchy, and it is these prediction signals that are modulated by attention. For example, the model proposed by Spratling [12] simulates the common physiological finding that attention to a stimulus enhances the firing rate of neurons tuned to specific stimulus features (e.g., orientation or colour for visual neurons), and has been shown to be mathematically equivalent to the biased competition model of attention [17–20]. In line with these alternative models, we investigated a second hypothesis – here termed the *feature information model* – which proposes that the interaction between attention and prediction at the level of neural responses is driven by changes in feature-specific information in the brain.

Here we tested whether the *feature information model* or the *mismatch information model* provides a better account of the neural coding of surprising stimuli in the human brain, and examined the influence of selective attention on each of these two neural codes. Participants attended to, or ignored, periodic streams of visual gratings, the orientations of which were either predictable, surprising, or unpredictable. We applied forward encoding models to whole-brain neural activity measured using EEG to quantify the neural selectivity for information related to the grating orientation and the mismatch between the predicted and observed grating orientations. We show that surprising stimuli evoke neural responses that contain information related to the difference between predicted and observed stimulus features, consistent with the *mismatch information model*. Crucially, we also find that attention increases the selectivity for mismatch information in the neural response to surprising stimuli, supporting the hypothesis that attention increases the gain of prediction errors [8].

## Results

We recorded brain activity using EEG as human observers (N = 24) undertook a rare-target detection task (see *Methods*; *Fig 1*). Participants fixated centrally and were presented with a periodic stream of gratings (100 ms duration, 500 ms ISI, 415 gratings per block) in one of two conditions (randomised across blocks). In *roving standard* blocks [21] (see *Fig 1A*), grating orientation was repeated between 4 and 11 times (*standards*) before changing to a new orientation (*deviants*, pseudo-randomly selected from one of nine orientations, spanning 0 - 160° in 20° steps). Grating orientation was thus ‘predictable’ for standards and ‘surprising’ for deviants. In *equiprobable blocks* [22] (see *Fig 1B*), gratings changed orientation on every presentation and thus could not be predicted (‘unpredictable’ *controls*). Attention was manipulated by having participants either monitor the grating stimuli for rare targets with a different spatial frequency (‘grating task’, *attended*), or ignore the gratings and instead monitor for rare fixation-dot targets with decreased contrast (‘dot task’, gratings *ignored*).

**Fig 1.**
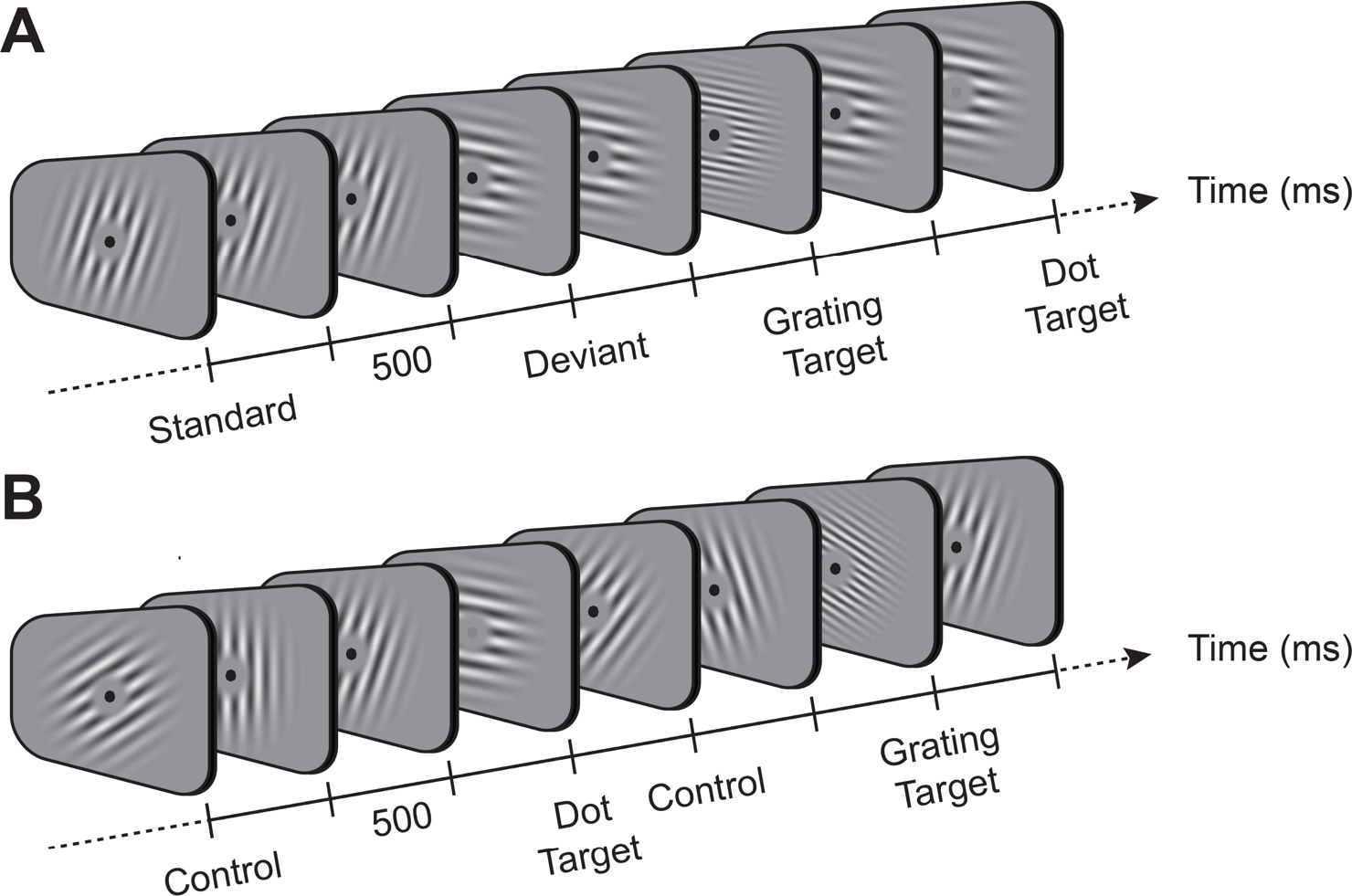
Example stimuli in each of the two block types used in the study. (A) Roving oddball sequence. In this sequence, the orientation of gratings was repeated over short sequences of stimuli (*standards*), before changing to a different orientation (*deviant*). During the grating or dot task, participants responded to rare gratings with high spatial frequency (*grating target*) or to rare decreases in fixation-dot contrast (*dot target*), respectively. (B) Equiprobable sequence. In this sequence, the orientation of control gratings changed with each successive presentation.

Participants completed the grating task and dot task in separate sessions, approximately one week apart (session order counterbalanced). At the beginning of each session, participants completed three practice blocks of the specified task, during which target salience levels were titrated to approximate a target detection rate of 75% (see *Methods*). Participants were then fitted with a 64-electrode EEG cap before completing 21 test blocks. One participant detected fewer than 50% of targets in both tasks and was therefore excluded from all further analyses. The remaining participants detected an equivalent percentage of targets in the grating task (75.64 ± 1.76%, mean ± SEM) and dot task (72.73 ± 2.54%; *t*(22) = 1.57, *p* = 0.13, *BF*_*10*_ = 0.12), and also produced similar numbers of false alarms in each (20.43 ± 3.79 and 22.57 ± 5.47, respectively; *t*(22) = −0.41, *p* = .684, *BF*_*10*_ = 0.18), suggesting that difficulty was well matched between attention conditions.

EEG data were pre-processed offline using EEGlab [23] and epoched according to the onset of each grating (see *Methods* for details). Statistical analyses were conducted using cluster-based permutation tests in Fieldtrip [24]. *S1 Fig* shows the main effects and interactions for the factors of attention and prediction on event related potentials (ERPs). Briefly, ERPs were modulated by both attention (86 - 434 ms, cluster-corrected *p* < .001; *S1A and S1C Figs*) and prediction (39 - 550 ms, cluster-corrected *p* < .001, *S1A Fig*). Follow-up analyses of the simple effects of prediction revealed that deviants elicited larger responses than both standards (39 - 550 ms, cluster-corrected *p* < .001; *S1A and S1D Figs*) and controls (324 - 550 ms, cluster-corrected *p* = .002; *S1A and S1E Figs*). The difference between deviants and controls emerged later and was smaller than the difference between deviants and standards, consistent with the notion that the former comparison reflects the pure effects of prediction (“genuine” mismatch response (MMR), [22]), whereas the latter comparison confounds the effects of prediction with those of adaptation to the standard (‘classic’ MMR, see [4] for a review).

We also observed an interaction between attention and prediction (180 - 484 ms, cluster-corrected *p* < .001; *S1A Fig*). Follow-up analyses revealed that attention increased both the classic MMR (176 - 469 ms, cluster-corrected *p* < .001; *S1F and S1G Figs*) and the genuine MMR (176 - 550 ms, cluster-corrected *p* < .001; *S1H and S1I Figs*). In the attended condition, both the classic and the genuine MMRs emerged approximately 200 ms after stimulus onset over posterior-lateral (PO7, PO8) electrodes (*S1B Fig*, solid green and yellow lines, respectively). Whereas the onset of the genuine MMR is consistent with previous literature [22], the classic MMR we report here emerged slightly later than what has typically been reported previously (∼150 ms; for a review see [4]). We note, however, that at least one previous study reported a visual MMR beginning as late as 250 ms [25], highlighting the variable nature of this component.

In the ignored condition, we observed classic and genuine MMRs (*S1B Fig*, dotted green and yellow lines, respectively) with positive polarities over posterior (PO7, PO8) and frontal (Fz) electrodes, respectively. In contrast, previous studies have typically (but not always; see [5]) reported mismatch *negativities*, even in the absence of attention [4]. A number of differences between previous studies and our own could explain this discrepancy (e.g. stimuli, interstimulus interval, presentation duration, task etc). In particular, we used large sinusoidal gratings (11° of visual angle) to optimise orientation decoding, in contrast to previous studies that presented much smaller oriented bars (∼3-4° of visual angle, e.g. [22,26]). Thus, the stimuli in the current study likely activated a larger area of visual cortex than those used in previous studies, which produced a different dipole (or combination of multiple dipoles) and associated projection to scalp electrodes (due to the complex folding structure of the cortex, [4]) than has previously been observed. Indeed, close inspection of the ERPs seems to indicate the presence of a single dipole projecting to frontal and posterior electrodes (note the highly similar pattern of activity between electrodes Fz and Pz, but with opposite sign, *S1A Fig*), which has not typically been observed in previous studies (e.g., note the relatively uniform responses across the scalp in [22,27,28]).

### Orientation information is enhanced with attention but not surprise

The *feature information model* predicts that the orientation-selective neural response to surprising stimuli (deviants) will be different to that of control stimuli. To investigate this hypothesis, we used a forward encoding model to estimate orientation selectivity from neural activity measured with EEG (see *Methods* for details). Briefly, we used multivariate regression to transform activity in electrode space into an orientation-selective ‘feature space’ [29–32], comprised of nine hypothetical ‘orientation channels’ matching those presented in the experiment (0 - 160°, in 20° steps). For each orientation channel, we modelled the expected activation across trials by convolving the presented orientation with a canonical orientation-selective tuning function. We then regressed this pattern of expected activity against the EEG data, separately for each time point (−100 - 550 ms after stimulus onset), to produce a weight matrix that converted multivariate activity in electrode space into activity in the specified orientation channel. The spatial weights for each orientation channel were then inverted to reconstruct the forward model and applied to an independent set of test trials (using a cross-validation procedure) to estimate activity across all orientation channels. As shown in *Fig 2A*, using the forward encoding approach we reconstructed distinct response profiles for each of the nine grating orientations presented to participants. Orientation channels were then realigned for each trial such that the presented orientation channel was centred on 0°, and activation patterns were averaged across trials in each condition. The forward encoding model revealed an orientation-tuned response throughout the epoch (*Fig 2B and 2C*). This response emerged soon after stimulus onset, peaked at ∼130 ms, and declined gradually until the end of the epoch.

**Fig 2.**
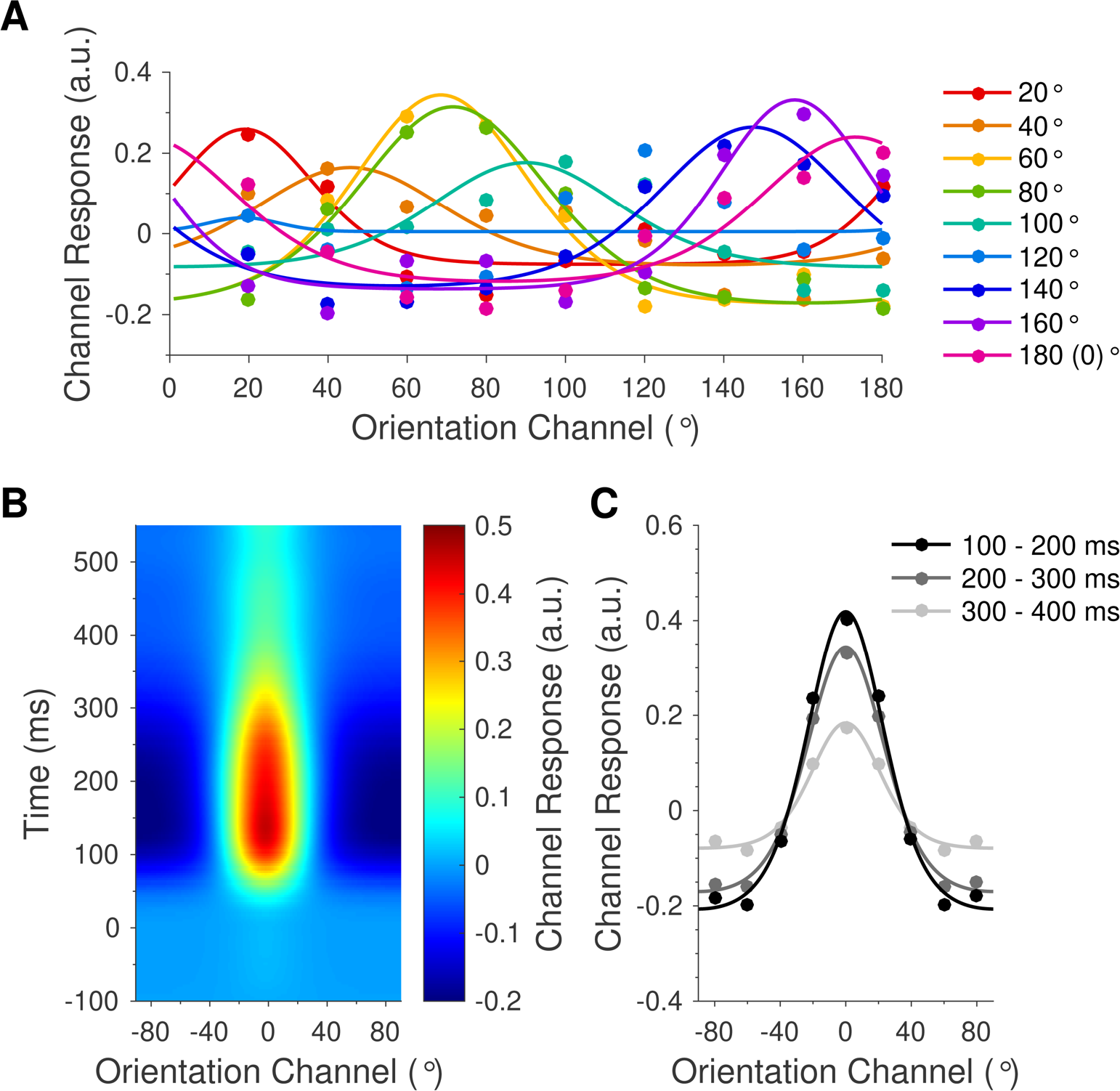
Stimulus-evoked orientation channel response profiles. **(A)** Reconstructed orientation channels, corresponding to each of the nine grating orientations presented to participants (0 - 160o, in 20o steps). Coloured dots indicate the modelled orientation channel activity across trials in which the labelled orientation was presented. Curved lines show functions fitted to the grand average data for illustrative purposes. Note that each coloured line is approximately centred on the presented orientation. **(B)** Time-resolved orientation response profile, centred on the presented orientation in each trial and averaged across participants and conditions. Orientation response profiles emerged shortly after stimulus onset and lasted until the end of the epoch. **(C)** Orientation response profiles, averaged across all participants and conditions in each of three successive 100 ms time windows. Dots show activation in each of the nine modelled orientation channels (mean-centred). Curved lines show functions fitted to the grand average data for illustrative purposes. Orientation information (response profile amplitude) was strongest from 100 – 200 ms and decreased throughout the epoch. Data are available at https://doi.org/10.17605/osf.io/a3pfq. a.u. = arbitrary units.

To quantify the effects of attention and prediction on orientation response profiles, we fitted the condition-averaged orientation channel responses with an exponentiated cosine function [33,34] using least squares regression:

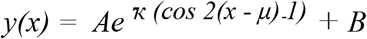

where *y* is the predicted orientation channel activity in response to a grating with orientation *x*; *A* is the peak response amplitude, *κ* is the concentration (i.e. inverse dispersion; a larger value corresponds to a “tighter” function), *μ* is the centre of the function, and *B* is the baseline offset (see *Methods*).

Attention increased the amplitude of orientation response profiles (219 - 550 ms, cluster-corrected *p* < .001; *Fig 3A and 3B*) but did not modulate the tuning concentration (all clusters *p* > .104). There was a significant main effect of prediction on the amplitude of orientation response profiles late in the epoch (324 – 550 ms, cluster-corrected *p* < .001; *S2C and S2D Figs*), as well as a non-significant but trending cluster early in the epoch (94 - 145 ms, cluster-corrected *p* = .154; *S2C Fig*, cluster not shown). Follow-up analyses revealed that orientation response profiles evoked by standards (0.11 ± 0.01 a.u.) were smaller than those of both deviants (0.25 ± 0.03 a.u.; *t*(22) = −4.32, *p* < 0.001, *BF*_*10*_ = 1469.10) and controls (0.22 ± 0.03 a.u.; *t*(22) = −3.79, *p* < 0.001, *BF*_*10*_ = 156.16; *S2C and S2D Figs*). Crucially, the amplitudes of orientation response profiles evoked by deviants and controls were equivalent (*t*(22) = 0.78, *p* = 0.443, *BF*_*10*_ = 0.19; *Fig 3A*, *S2C and S2D Figs*). Finally, there was no effect of prediction on the concentration of orientation response profiles (all clusters *p* > .403), and no interaction between attention and prediction on either the amplitude (cluster-corrected *p* = .093, *S2E and S2F Figs*) or concentration (no clusters found) of orientation response profiles.

**Fig 3.**
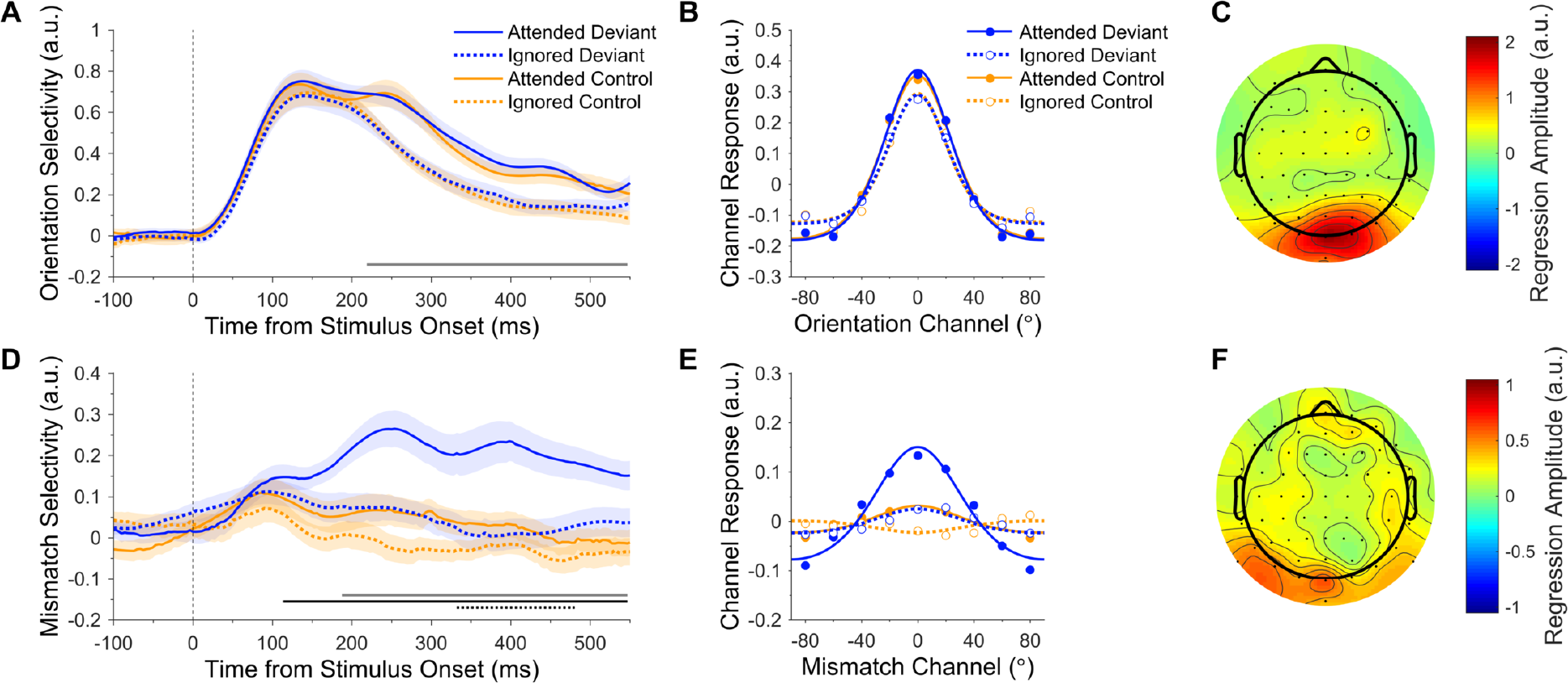
Effects of attention and prediction error on orientation and mismatch response profiles. **(A-C)** Orientation response profiles. (**A**) Orientation selectivity (response profile amplitude) for each condition over time. Shading indicates the SEM. Thin black lines indicate differences between deviants and controls, separately for attended and ignored stimuli. The dark grey bar along the x-axis indicates the main effect of attention (cluster-corrected). (**B**) Orientation response profiles, averaged across the significant effect of attention shown in **A**(219 - 550 ms). Dots show activation in each of the nine modelled mismatch channels. Curved lines show functions fitted to channel responses (fitted to grand average data for illustrative purposes). (**C**) Univariate sensitivity for stimulus orientation across all conditions (see *Methods*). Topography shows the permutation-corrected z-scores, averaged across the significant effect of attention shown in **A** (219 - 550 ms). Posterior electrodes were the most sensitive to orientation information. **(D-F)** Mismatch response profiles (observed minus predicted orientation). **(D)** Mismatch selectivity (response profile amplitude) for each condition over time. The grey, solid black, and dotted black bars along the x-axis indicate the main effect of attention, main effect of prediction, and the interaction, respectively (cluster-corrected). Attention enhanced the mismatch response profile in response to deviants but not controls. (**E**) Mismatch response profiles, collapsed across the significant interaction shown in **D** (332 – 480 ms). (**F**) Univariate sensitivity for mismatch response profiles evoked by attended deviants (see *Methods*), averaged across 332 – 480 ms. Posterior electrodes were the most sensitive to mismatch information. Note that **C** and **F** use different scales. Data are available at https://doi.org/10.17605/osf.io/a3pfq. a.u. = arbitrary units.

To determine the scalp topography that was most informative for orientation encoding, we calculated univariate sensitivity separately for each electrode across all trials, and averaged across time points in the significant main effect of attention (see *Methods*). As revealed in *Fig 3C*, posterior electrodes were the most sensitive to orientation information, as would be expected for a source in visual cortex.

### Attention facilitates the neural encoding of mismatch information

The *mismatch information model* proposes that prediction errors are represented in populations of neurons tuned to the *difference* between predicted and observed stimulus features. According to this model, therefore, surprising stimuli (deviants) should produce a more mismatch-selective neural response than control stimuli. Furthermore, if attention enhances the gain of prediction errors [8], we should expect an interaction between attention and prediction, such that attention enhances the amplitude of mismatch response profiles evoked by deviants more than that of controls, because deviants should evoke a larger prediction error [2]. To investigate these hypotheses, we trained a separate forward encoding model, as described above, on the angular difference between gratings (deviants or controls) and the preceding stimuli. That is, deviants were coded according to the difference between the deviant orientation and the preceding standard orientation, and controls were coded according to the difference between successive control orientations. For example, if a horizontally oriented deviant (0°) was preceded by a standard that was oriented at 40° (clockwise of horizontal), it would be coded as a mismatch of −40° (0 - 40°).

As shown in *Fig 3D and 3E*, we were able to reconstruct mismatch response profiles for attended deviants. By contrast, mismatch response profiles were clearly weaker in response to controls and ignored deviants. There was a significant main effect of attention on the amplitude of mismatch response profiles (attended > ignored, 188 – 550 ms, cluster-corrected *p* = .002; *Fig 3D*, grey bar along x-axis). There was also a significant main effect of prediction (deviant > control, 113 – 550 ms, cluster-corrected *p* < .001; *Fig 3D*, solid black bar along x-axis), suggesting that prediction error is encoded according to the mismatch between predicted and observed features. Crucially, attention and prediction interacted to influence the amplitude of mismatch response profiles (332 – 480 ms, cluster-corrected *p* = .031; *Fig 3D*, dotted black bar along x-axis). As can be seen in *Fig 3D and 3E*, attention enhanced the amplitude of deviant mismatch response profiles but had little effect on those evoked by controls, supporting the hypothesis that attention boosts prediction errors [8].

The concentration of mismatch response profiles was not modulated by attention (all clusters *p* > .888) or the interaction between attention and prediction (all clusters *p* > .615), although we did find a significant main effect of prediction on the concentration of mismatch response profile fits (controls > deviants, 344 - 422 ms, cluster-corrected *p* < .001). Since controls seemed to produce negligible mismatch response profiles during this time period (yellow lines, *Fig 3D*), however, we followed up this result by averaging mismatch response amplitudes across the significant timepoints and comparing these values to zero with a *t*-test and Bayes Factor analysis (uniform prior, lower bound: 0, upper bound = 0.3). We found that control mismatch response profile amplitudes (.005 ± .023 a.u.) were equivalent to zero (*t*(22) = 0.19, *p* = .848, *BF*_*10*_ = 0.11), suggesting that the observed effect on concentration was more likely an artefact of the fitting procedure than a true effect of prediction on mismatch response profiles.

We calculated the sensitivity of each electrode to mismatch information in trials that contained attended deviants, and collapsed across the significant interaction between 332 and 480 ms. As revealed in *Fig 3F*, posterior electrodes were again the most informative, but the topography of mismatch sensitivity was weaker and more sparsely distributed than that of orientation decoding (*Fig 3C*).

### Mismatch information increases with the strength of predictions

Next, we investigated whether the number of preceding standards was related to the amplitude of prediction error response profiles. Repeated presentations of the standard are thought to increase the strength of the memory trace, resulting in larger prediction errors to a subsequent surprising stimulus [35]. Mismatch response profiles evoked by attended deviants were grouped according to the number of preceding standards (4-7 repetitions vs 8-11 repetitions) and fitted with exponentiated cosine functions (see *Methods*). As can be seen in *Fig 4A and 4B*, increasing the number of standard repetitions also increased the amplitude of mismatch response profiles (387 - 520 ms, cluster-corrected *p* = .050). This finding is consistent with the notion that successive standards allow a more precise prediction to be generated, which results in enhanced prediction errors when violated. Finally, there was no effect of the number of standard repetitions on the concentration of mismatch response profiles (cluster-corrected *p* = .314).

**Fig 4.**
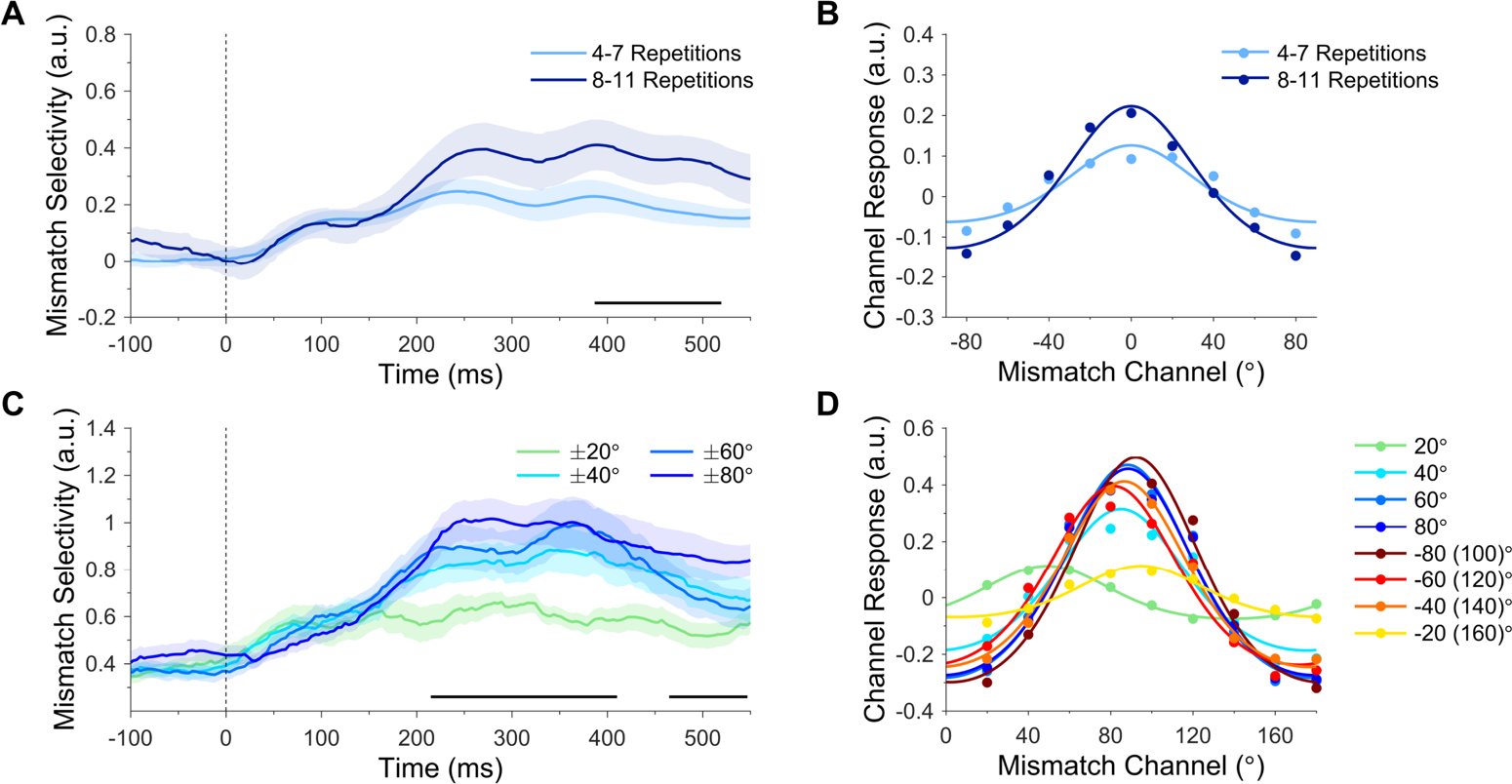
Mismatch response profiles (putative prediction error) evoked by attended deviants. (**A**) Effect of standard repetition on mismatch selectivity (response profile amplitude). Mismatch response profiles evoked by attended deviants were larger following long standard sequences (8 - 11 repetitions) than short standard sequences (4 - 7 repetitions). The black bar along the x-axis denotes significant differences (cluster-corrected). (**B**) Mismatch response profiles, collapsed across significant time points in **A** (387 - 520 ms). Dots show activation in each of the nine modelled mismatch channels. Curved lines show functions fitted to channel responses (fitted to grand average data for illustrative purposes). (**C**) Effect of deviation angle on mismatch selectivity. Mismatch response profile amplitude increased with the magnitude of deviation (±80° > ±20°). (**D**) Mismatch response profiles for each deviation angle, collapsed across the earlier cluster shown in **C** (215 - 410 ms). Curved lines show functions fitted with a variable centre (fitted to grand average data for illustrative purposes). Data are available at https://doi.org/10.17605/osf.io/a3pfq. a.u. = arbitrary units.

### Mismatch information increases with the magnitude of violation

We also tested whether larger deviations from the prediction increased selectivity for mismatch information. Mismatch response profiles of attended deviants were grouped according to the angular difference between the deviant and preceding standard (i.e., the original mismatch values entered into the encoding model) and fitted with exponentiated cosine functions (variable centre, see *Methods*). There was a significant main effect of deviation magnitude on mismatch response profile amplitude (215 - 410 ms, cluster-corrected *p* = .004). As shown in *Fig 4C*, the amplitude of mismatch response profiles increased with the absolute deviation angle (±80° > ±60° > ±40° > ±20°), supporting the notion that larger angular deviations (from the predicted orientation) produce more prediction error. A second cluster emerged later in the epoch (465 - 550 ms, cluster-corrected *p* = .031), which followed a similar pattern but with the amplitude of the ±40° and ±60° responses reversed. Intriguingly, individual mismatch response profiles were typically centred on the orthogonal deviation angle (90°, *Fig 4D*). This pattern of results differs from the individual orientation response profiles (*Fig 2A*), which were (approximately) centred on the presented orientation.

### Attention produces temporally stable mismatch response profiles

In a final step, we investigated whether the spatial maps that produce mismatch response profiles are stable or evolve dynamically over time. We used the same forward encoding analysis as above, with the exception that the trained weights at each time point were tested on *all* time points in the epoch [30,36] (see *Methods*). This produced a *train time* x *test time* generalisation matrix of mismatch channel responses, to which we fitted exponentiated cosine functions. *Fig 5* shows the mismatch selectivity (response profile amplitude) for attended and ignored deviants, generalised across time. As revealed in *Fig 5A*, the mismatch response profile evoked by attended deviants generalised across the latter part of the epoch (black outline surrounding large red patch in upper right quadrant between ∼200 - 550 ms, cluster-corrected *p* = .010), indicating that the spatial map associated with mismatch information was relatively consistent throughout this period. Note also that this pattern of generalisation was asymmetrical (triangular-shaped, rather than square-shaped). Specifically, the spatial map trained at ∼450 ms generalised to the (test) time point at ∼250 ms, but training at ∼250 ms did not generalise to testing at ∼450 ms. Since asymmetrical generalisation can indicate differences in signal-to-noise ratios between time points [36], this finding suggests that the strength of prediction error signals may have increased toward the end of the epoch. It is also worth noting that the apparent generalisation of spatial maps trained at stimulus onset (t_train_ = 0) to later times in the epoch (∼200 – 550 ms, red patch along the x-axis) was not significantly different from zero (no clusters found in this region) and produced high residuals in the function fits (see *S3 Fig*), suggesting that this pattern represents noise. Finally, the mismatch response profile evoked by ignored stimuli (*Fig 5B*) did not generalise across time points (all clusters *p* > .935) and was significantly smaller than that of attended stimuli (significant difference denoted by the opaque patch in *Fig 5C*; *p* = .026).

**Fig 5.**
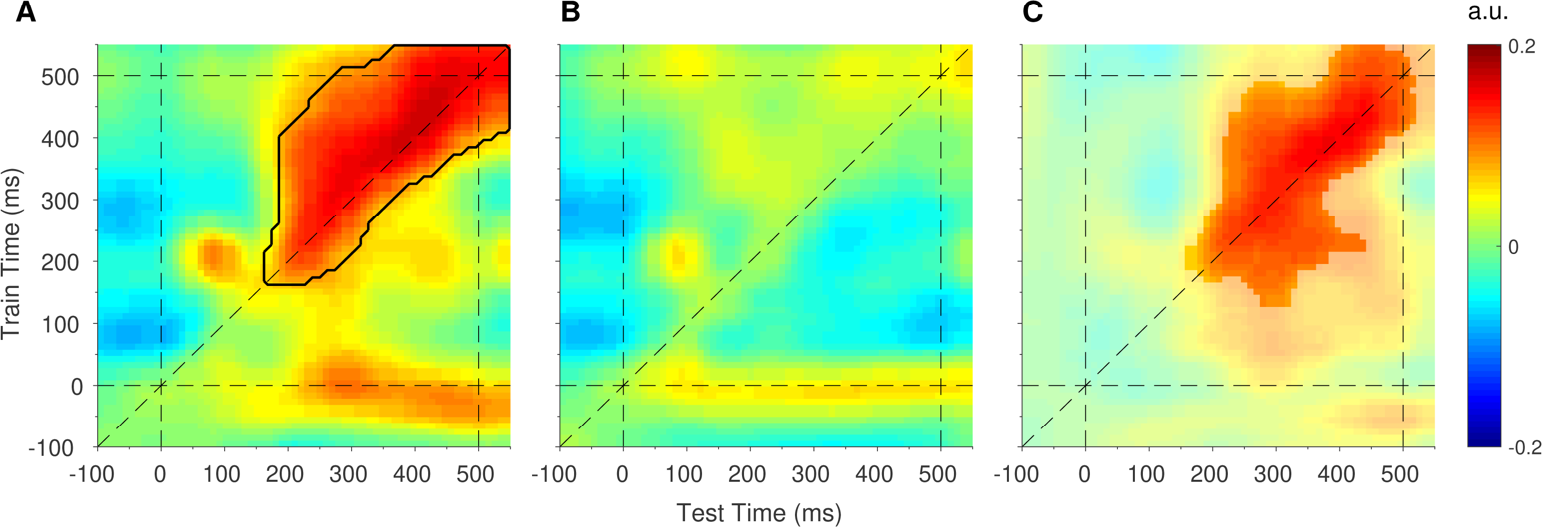
Generalised mismatch response profiles in response to (**A**) attended deviants and (**B**) ignored deviants. The dashed diagonal line indicates on-axis encoding (equivalent to the time-series plot in *Fig 4A*). The black outline shows mismatch response profiles significantly larger than zero (cluster-corrected). (**C**) Difference map (attended minus ignored), thresholded to show the significant effect of attention on mismatch response profiles (cluster-corrected). Data are available at https://doi.org/10.17605/osf.io/a3pfq. a.u. = arbitrary units.

## Discussion

Here we set out to determine what type of information is modulated in the interaction between attention and prediction [8]. To achieve this, we used forward encoding models of EEG data to quantify the selectivity for orientation and mismatch information in the neural responses to surprising and unpredictable stimuli in the well-established roving oddball paradigm [21,37]. Relative to unpredictable stimuli (controls), we found that EEG responses to surprising stimuli (deviants) were equally selective for orientation information, but more selective for information related to the *difference* between predicted and observed stimulus features. These results are consistent with the *mismatch information model*, and support the idea that top-down prediction signals ‘cancel out’ matching bottom-up sensory signals and leave only the remaining prediction error to propagate forward [2,3,8,9]. Crucially, we also found that attention increased the selectivity for mismatch information in neural responses to surprising but not control stimuli. This finding demonstrates that attention boosts mismatch information evoked by surprising stimuli (putative prediction errors), and is consistent with a recent version of predictive coding theory that proposes attention optimises the expected precision of predictions by increasing the gain of prediction errors [8].

We found no difference between orientation response profiles evoked by surprising and unpredictable stimuli (a prediction of the *feature information model*), suggesting that the increase in EEG activity that is typically observed with surprise is not coded according to stimulus features. This finding contradicts predictive coding models in which predictions (or ‘representations’) of stimulus features are passed up the visual hierarchy [12,16,17]. Because feedforward connections largely originate primarily from superficial pyramidal cells and it is this activity that is measured with EEG [9,14,15], these models would predict that surprise changes the feature-selectivity of EEG responses: a finding we do not observe here. This finding might also seem to contradict a recent study that demonstrated greater selectivity for orientation information in early visual cortex BOLD activity following presentation of a predicted grating, relative to a surprising grating [38]. Since BOLD activity indirectly measures the activity patterns of heterogenous populations of neurons, however, this change in feature-selectivity could have reflected a change in either of the two neuronal populations proposed to underlie predictive coding - *predictions* or *prediction errors*. The latter interpretation is inconsistent with the results of the present study, which suggests that prediction errors are encoded according to the mismatch between predicted and observed stimulus features, and not the features themselves. The former interpretation (i.e. that predictions are coded according to the stimulus features) fits well with a recent study that showed prediction induces feature-specific templates immediately prior to stimulus onset [31]. Thus, a parsimonious account of the literature to date suggests that *predictions* and *prediction errors* are represented in the brain via distinct neural codes: whereas predictions are represented according to stimulus features, prediction errors are represented according to the *mismatch* between predicted and observed stimulus features.

In a recent study by our group [39], we observed a decrease in orientation selectivity in the neural response to predicted stimuli, relative to surprising stimuli, shortly after stimulus onset (79 – 185 ms). Here we observed a similar (but non-significant) trend in the same direction (standards < deviants) at approximately the same time (94 - 145 ms, *S2C Fig*, cluster not shown). Close inspection of the present results, however, suggests that some orientation information evoked by the previous standard was still present in the brain at the onset of the subsequent standard (indicated by the above-zero amplitude of the orientation response to standards at stimulus onset, t = 0 ms, *S2C Fig*), which may have obscured detection of the early effect reported in Tang et al. [39]. Interestingly, the present results revealed a late effect of prediction (standards < deviants, 324-550 ms, *S2C and S2D Figs*) that was not observed in our previous work [39]. Since a critical difference between the two studies was the number of times identical stimuli could be presented consecutively (no more than twice in the previous study), we speculate that the late effect observed here might reflect the minimal amount of model-updating required after the presentation of a precisely predicted stimulus.

We also found that attention increased the amplitude of orientation response profiles (*Fig 3A and 3B*), consistent with previous studies that applied forward encoding models to human fMRI [34,40] and time-frequency-resolved EEG data [29]. The present study replicates and extends these studies with the application of forward encoding models to time-resolved EEG recordings (resulting in <30 ms temporal resolution after smoothing), demonstrating that attention increases feature selectivity in the human brain from approximately 200 ms after stimulus onset.

Crucially, we also tested the interactive effects of attention and prediction on information processing in the brain. There was a large and significant effect of attention on mismatch response profiles in response to surprising but not unpredictable stimuli (beginning around 150 ms after stimulus onset and reaching significance from ∼350 ms). This finding demonstrates that attention boosts prediction errors evoked by surprising stimuli, and is consistent with a recent iteration of predictive coding theory according to which attention optimises the expected precision of prediction errors [8]. Previous studies have found evidence for an interaction between attention and prediction in both the auditory [5] and visual [6,41] modalities. Importantly, these studies used *activation-based* analyses to compare differences between predicted and unpredicted stimuli at the level of overall neural activity, but did not investigate what *type of information* is modulated in the interaction between attention and prediction. In contrast, the present study used *information-based* analyses [42] to identify specific patterns of neural activity that are associated with orientation-mismatch information in the brain, and showed that selectivity for this type of information (but not feature information) is increased with attention. Thus, the present study provides clear support for the hypothesis that attention boosts the gain of prediction errors [8]. It will be important for future research to investigate whether the interactive effects of attention and prediction on mismatch information is contingent on the type of attention (e.g., feature-based versus spatial attention) or prediction (e.g., rule-based versus multimodal cue-stimulus predictions; [31,43]).

Interestingly, we found that the magnitude of mismatch response profiles correlated with the number of preceding standards (*Fig 4A and 4B*). Previous work in the auditory domain demonstrated that successive repetitions of the standard evoke progressively increased responses to a subsequent attended deviant [35]. Here we find a corollary for this effect in the visual domain and demonstrate that the neural activity modulated by the number of preceding standards is likely encoded as mismatch information. This finding is also consistent with the notion that repeating the standard allows a more precise prediction to be generated, which results in a larger prediction error to a subsequent surprising stimulus [44].

We also found that mismatch response profiles increased with the magnitude of the mismatch between predicted and observed stimulus features (*Fig 4C*). Previous work in the auditory domain has demonstrated a correlation between deviation magnitude and the amplitude of the neural response to deviants (i.e. the mismatch negativity) [45]. Here we demonstrate a relationship between deviation magnitude and selectivity for mismatch information (as opposed to activation levels) in the visual domain, suggesting that the magnitude of mismatch information might be used by the brain to guide updating of the predictive model. Since the present study investigated mismatch signals with respect to a continuous and circular feature dimension (i.e. orientation), it will be important for future research to extend the current line of research to non-circular (e.g. luminance, auditory frequency) and categorical (e.g. facial emotions) feature dimensions.

Somewhat surprisingly, there was a lateral shift in the response profile of individual mismatch channels toward the orthogonal (90°) channel (*Fig 4D*). The extent of this effect depended on the deviation magnitude, with large deviations (±40-80°) being predominantly stacked over the 90° channel and smaller deviations (±20°) being more closely aligned with their veridical mismatch angle (*Fig 4D*). We speculate that this might indicate a qualitative difference in the way that small and large prediction errors were treated by the brain in the present study. Small deviations may have resulted in updating and retention of the current model (via a near-veridical mismatch signal), whereas large deviations may have resulted in the wholesale rejection of the current model (via a generic mismatch signal) in favour of an alternative model that represents the deviant stimulus. In the latter case, the magnitude of the (orthogonal) mismatch channel response might represent an efficient code that the brain utilises to select from a number of likely alternative models.

Intriguingly, a number of recent studies failed to find an interaction between the effects of attention and prediction on stimulus information in the brain [31,38,46]. If predictions are encoded according to stimulus features, as we argue above, these null findings contradict the theory that attention boosts predictions [47]. In contrast, we show that *prediction errors*, represented according to the *mismatch* between predicted and observed stimulus features, are enhanced with attention. Although the present study cannot speak to the activity of single neurons, we note that the emerging picture is consistent with the notion that predictions and prediction errors are represented in distinct populations of neurons [2] that encode two distinct types of information and are differentially influenced by attention. Under this framework, feature information encoded by prediction units would be immune to attention, whereas mismatch information encoded by *prediction error* units would be enhanced by attention. Future research could test these hypotheses at the single-cell level, for example by using single-unit electrode recordings or 2-photon calcium imaging to assess whether different neurons within a given cortical area satisfy these constraints.

## Methods

### Ethics Statement

The study was approved by The University of Queensland Human Research Ethics Committee (approval number: 2015001576) and was conducted in accordance with the Declaration of Helsinki. Participants provided informed written consent prior to commencement of the study.

### Participants

Twenty-four healthy participants (11 female, 13 male, mean = 23.25 years, SD = 9.01 years, range: 18 to 64 years) with normal or corrected-to-normal vision were recruited via an online research participation scheme at The University of Queensland.

### Stimuli

Stimuli were presented on a 61 cm LED monitor (Asus, VG248QE) with a 1920 × 1080 pixel resolution and refresh rate of 120 Hz, using the PsychToolbox presentation software [48] for Matlab (v.15b) running under Windows 7 with a NVidia Quadro K4000 graphics card. Participants were seated in a comfortable armchair in an electrically shielded laboratory, with the head supported by a chin rest at a viewing distance of 57 cm.

During each block, 415 gratings with Gaussian edges (outer diameter: 11°, inner mask diameter: 0.83°, spatial frequency: 2.73 c/°, 100% contrast) were presented centrally for 100 ms with a 500 ms ISI. Grating orientations were evenly spaced between 0° (horizontal) and 160° (in 20° steps). Eighteen (18) gratings in each block (2 per orientation) were presented with a higher spatial frequency (range: 2.73 - 4.55 c/°, as per staircase procedure, below), with a gap of at least 1.5 s between any two such gratings. We used a modified de Bruijn sequence to balance the order of grating orientations across conditions, sessions, and participants. Specifically, we generated two 9-character (orientation) sequences without successive repetitions (e.g. ABCA, not ABCC) - one with a 3-character sub-sequence (504 characters long) and another with a 2-character sub-sequence (72 characters long) - and appended two copies of the former sequence to three copies of the latter sequence (1224 characters in total). This master sequence was used to allocate the order of both deviants and controls in each session (using different, random start-points), and ensured that each orientation was preceded by equal numbers of all other orientations (up to 2+ preceding stimuli) so that decoding of any specific orientation could not be biased by the orientation of preceding stimuli.

In roving oddball sequences, the number of Gabor repetitions (i.e., standards) was balanced across orientations within each session, such that each orientation repeated between 4 and 11 times according to the following distribution: (31, 31, 31, 23, 5, 5, 5, 5), respectively. During each block, the fixation dot (diameter: 0.3°, 100% contrast) decreased in contrast 18 times (contrast range: 53-98% as per staircase procedure, below) for 0.5 s (0.25 s linear ramp on and off). Contrast decrement onsets were randomised separately for each block, with a gap of at least 1.5 s between any two decrement onsets.

### Procedure

Participants attended two testing sessions of 60 minutes duration, approximately one week apart, and completed one of two tasks in each session (*Fig 1*, session order counterbalanced across participants). For the grating task, participants were informed that approximately 1/20 of the gratings would be a target grating with a higher spatial frequency than non-targets, and were asked to press a mouse button as quickly as possible when they detected a target grating; all other gratings were to be ignored. For the dot task, participants were informed that the fixation dot would occasionally decrease in contrast, and were asked to press a mouse button as quickly as possible when they detected such a change. Participants initially completed three practice blocks (3.5 min per block) with auditory feedback (high or low tones) indicating missed targets and the accuracy of their responses. During practice blocks in the first testing session, target salience (spatial frequency or dot contrast change, depending on the task) was adjusted dynamically using a Quest staircase procedure [49] to approximate 75% target detection. During practice blocks in the second testing session, target salience was adjusted to approximate the same level of target detection observed in the first testing session. Participants were requested to minimise their number of false alarms. After the practice blocks, participants were fitted with an EEG cap (see *EEG Data Acquisition*) before completing a total of 21 test blocks (3 equiprobable, 18 roving standard, block order randomised) without auditory feedback. After each block participants were shown the percentage of targets correctly detected, the speed of these responses, and how many non-targets were responded to (false alarms).

### Behavioural Data Analysis

Participant responses were scored as hits if they occurred within one second of the onset of a target grating in the grating task, or within one second of the peak contrast decrement in the dot task. Target detection was then expressed as a percentage of the total number of targets presented in each testing session. One participant detected less than 50% of targets in both sessions and was removed from further analysis. Target detections and false alarms across the two sessions were compared with paired-samples *t*-tests and Bayes Factors. Bayes factors allow for quantification of evidence in favour of either the null or alternative hypothesis, with *B*_*01*_ > 3 indicating substantial support for the alternative hypothesis and *B*_*01*_ < 0.33 indicating substantial support for the null hypothesis [50]. Bayes factors were computed using the Dienes [50,51] calculator in Matlab, with uniform priors for target detection (lower bound: −25%, upper bound: 25%) and false alarms (lower bound: −50, upper bound: 50).

### EEG Data Acquisition

Participants were fitted with a 64 Ag-AgCl electrode EEG system (BioSemi Active Two: Amsterdam, Netherlands). Continuous data were recorded using BioSemi ActiView software (http://www.biosemi.com), and were digitized at a sample rate of 1024 Hz with 24-bit A/D conversion and a .01 – 208 Hz amplifier band pass. All scalp electrode offsets were adjusted to below 20μV prior to beginning the recording. Pairs of flat Ag-AgCl electro-oculographic electrodes were placed on the outside of both eyes, and above and below the left eye, to record horizontal and vertical eye movements, respectively.

### EEG Data Preprocessing

EEG recordings were processed offline using the EEGlab toolbox in Matlab [23]. Data were resampled to 256 Hz and high-pass filtered with a passband edge at 0.5 Hz (1691-point Hamming window, cut-off frequency: 0.25 Hz, −6 db). Raw data were inspected for the presence of faulty scalp electrodes (2 electrodes, across 2 sessions), which were interpolated using the average of the neighbouring activations (neighbours defined according to the EEGlab Biosemi 64 template). Data were re-referenced to the average of all scalp electrodes, and line noise at 50 and 100 Hz was removed using the Cleanline plugin for EEGlab (https://www.nitrc.org/projects/cleanline). Continuous data were visually inspected and periods of noise (e.g., muscle activity) were removed (1.4% of data removed in this way, across sessions).

For artefact identification, the cleaned data were segmented into 500 ms epochs surrounding grating onsets (100 ms pre- and 400 ms post-stimulus). Improbable epochs were removed using a probability test (6SD for individual electrode channels, 2SD for all electrode channels, 6.5% of trials across sessions), and the remaining data were subjected to independent components analyses (ICA) with a reduced rank in cases of a missing EOG electrode (2 sessions) or an interpolated scalp electrode (2 sessions). Components representing blinks, saccades, and muscle artefacts were identified using the SASICA plugin for EEGlab [52].

For further analysis, the cleaned data (i.e., prior to the ICA analysis) were segmented into 800 ms epochs surrounding grating onsets (150 ms pre- and 650 ms post-stimulus). Independent component weights from the artefact identification process were applied to this new data set, and previously identified artefactual components were removed. Baseline activity in the 100 ms prior to each stimulus was removed from each epoch. Grating epochs were then separated into their respective attention and prediction conditions. Epochs in the grating task were labelled as ‘Attended’ and epochs in the dot task were labelled as ‘Ignored’. Epochs in the roving oddball sequence were labelled as ‘Deviants’ when they contained the first stimulus in a repeated train of gratings, and ‘Standards’ when they contained a grating that had been repeated between five and seven times. Epochs in the equiprobable sequence were labelled as ‘Controls’.

### Event-Related Potential Analyses

Trials in each attention and prediction condition were averaged within participants to produce event-related potentials (ERPs) for each individual. The effect of attention was assessed using a two-tailed cluster-based permutation test across participant ERPs (Monte-Carlo distribution with 5000 permutations, *p*_*cluster*_<0.05; sample statistic: dependent samples *t*-statistic, aggregated using the maximum sum of significant adjacent samples, *p*_*sample*_<.05). Because there were three, rather than two, levels of prediction, we tested the effect of prediction with a cluster-based permutation test that used *f*-statistics at the sample level and a one-sided distribution to account for the positive range of *f*-statistics (Monte-Carlo distribution with 5000 permutations, *p*_*cluster*_<0.05; sample statistic: dependent samples *f*-statistic, aggregated using the maximum sum of significant adjacent samples, *p*_*sample*_<.05). Simple contrasts between prediction conditions (deviants vs standards, and deviants vs controls) were tested using two-tailed cluster-based permutation tests (with the same settings as used to investigate attention). The interaction between attention and prediction was assessed by subtracting the ignored ERP from the attended ERP within each prediction condition and subjecting the resulting difference waves to a one-tailed cluster-based permutation test across participant ERPs (Monte-Carlo distribution with 5000 permutations, *p*_*cluster*_<0.05; sample statistic: dependent samples *f*-statistic, aggregated using the maximum sum of significant adjacent samples, *p*_*sample*_<.05). The interaction effect was followed-up by comparing difference waves (attended - ignored) between deviants and standards, and between deviants and controls (two-tailed cluster-based permutation tests, same settings as above).

### Forward Encoding Models

To investigate the informational content of orientation signals, we used a forward encoding model [29,53] designed to control for noise covariance in highly correlated data [31,54; https://github.com/Pim-Mostert/decoding-toolbox], such as EEG. We modelled an idealised basis set of the nine orientations of interest (0-160° in 20° steps) with nine half-wave rectified cosine functions raised to the 8^th^ power, such that the response profile associated with any particular orientation in the 180° space could be equally expressed as a weighted sum of the nine modelled orientation channels [29]. We created a matrix of nine regressors that represented the grating orientation presented on each trial in the training set (1 = the presented orientation, 0 = otherwise) and convolved this regressor matrix with the basis set to produce a design matrix, ***C*** (9 orientation channels x *n* trials). The EEG data could thus be described by the linear model:

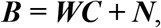

where ***B*** represents the data (64 electrodes x *n* trials), ***W*** represents a spatial weight matrix that converts activity in channel space to activity in electrode space (64 electrodes x 9 orientation channels) and ***N*** represents the residuals (i.e., noise).

To train and test the forward encoding model, we used a three-fold cross-validation procedure that was iterated 100 times to increase reliability of the results. Within each cross-validation iteration, the experimental blocks were folded into thirds: one third of trials served as the test set and the remaining two-thirds served as the training set, and folds were looped through until each fold had served as a test set. Across successive iterations of the cross-validation procedure, the number of trials in each condition was balanced within folds by random selection (on the first iteration) or by selecting the trials that had been utilised the least across previous folds (subsequent iterations).

Prior to estimating the forward encoding model, each electrode in the training data was demeaned across trials, and each time point was averaged across a 27.3 ms window centred on the time point of interest (corresponding to an *a priori* window of 30 ms, rounded down to an odd number of samples to prevent asymmetric centring). Separately for each time point and orientation channel of interest, *i*, we solved the linear equation using least square regression:

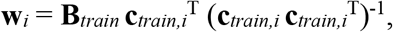

where **w**_*i*_ represents the spatial weights for channel *i*, **B**_*train*_ represents the training data (64 electrodes x n_*train*_ trials), and **c**_*train*,*i*_ represents the hypothetical response of channel *i* across the training trials (1 x n_*train*_ trials). Following Mostert et al. [54], we then derived the optimal spatial filter **v**_*i*_ to recover the activity of the *i*th orientation channel:

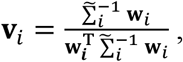

where Σ_*i*_ is the regularized covariance matrix for channel *i*, estimated as follows:

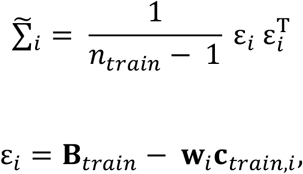

where *n*_*train*_ is the number of training trials. The covariance matrix 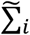 was regularized by using the analytically determined shrinkage parameter [31]. Combining the spatial filters across each of the nine orientation channels produced a channel filter matrix **V**(64 electrodes x 9 channels).

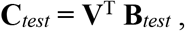

where **B**_*test*_ represents the test data at the time point of interest (64 electrodes x *n*_*test*_ trials), averaged over a 27.3 ms window (as per the training data). Finally, the orientation channel responses for each trial were circularly shifted to centre the presented orientation on 0°, and the zero-centred responses were averaged across trials within each condition to produce the condition-average orientation channel response (*Fig 3B*).

To assess information related to the mismatch between predicted and observed stimulus features (*Fig 3D and 3E*), we computed a second forward encoding model as above, with the exception that now the regression matrix represented the difference between the current grating orientation (deviant or control) and the previous grating orientation (standard or control, respectively). That is, a grating at 60° orientation that followed a grating at 20° orientation would be coded as 40° (current minus previous orientation).

To assess the dynamic nature of mismatch response profiles (*Fig 5*), we trained the weight matrix, *W*, at a single time point in the training set, *B*_1_ (using a 30 ms sliding window), and then applied the weights to *every third* time point in the test set, *B*_2_ (using a 30 ms sliding window). This process was repeated for every third time point in the training set, resulting in a 3-dimensional matrix that contained the population response profile at each cross-generalised time point (9 orientations x 66 training time points x 66 testing time points).

### Quantifying Channel Responses

Previous studies have utilised a number of different methods to quantify the selectivity of neural response profiles [30,31]. Since we were interested in characterising the properties of neural response profiles, we opted to fit an exponentiated cosine function to the modelled data [33,34] using least square regression:

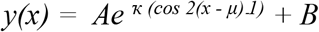

where *y* is the predicted orientation channel activity in response to a grating with orientation *x*; *A* is the peak response amplitude, *κ* is the concentration parameter, *μ* is the centre of the distribution, and *B* is the baseline offset. Fitting was performed using the non-linear least square method in MATLAB (trust region reflective algorithm). The free parameters *A*, *κ*, and *B* were constrained to the ranges −0.5, 2), (1.5, 200), and (−1.0, 0.5), respectively, and initiated with the values 0.5, 2, and 0, respectively. The free parameter *μ* was constrained to be zero when quantifying mean-centred orientation or mismatch response profiles (which should be centred on zero, *Figs 3, 4A and 4B*). When quantifying individual (uncentred) mismatch channel response profiles (*Fig 4C and 4D*), the free parameter *μ* was allowed to vary between −90° and 90°. To reduce the likelihood of spurious (inverted) fits, the parameter search was initiated with a *μ* value centred on the channel with the largest response.

The main effects of attention and prediction on orientation or mismatch response profiles were assessed with cluster-based permutation tests across participant parameters (amplitude, concentration). The interaction effects (between attention and prediction) on orientation and mismatch response profiles were assessed by first subtracting the ignored response from the attended response, and then subjecting the resulting difference maps to cluster-based permutation tests. In cases where two levels were compared (i.e. the main effect of attention on orientation response profiles, and all effects on mismatch response profiles), we used two-tailed cluster-based permutation tests across participant parameters (Monte-Carlo distribution with 5000 permutations, *p*_*cluster*_<0.05; sample statistic: dependent samples *t*-statistic, aggregated using the maximum sum of significant adjacent samples, *p*_*sample*_<.05). In cases where three levels were compared (i.e. the main effect of prediction and the interaction effect on orientation response profiles), we used one-tailed cluster-based permutation tests across participant parameters (Monte-Carlo distribution with 5000 permutations, *p*_*cluster*_<0.05; sample statistic: dependent samples *f*-statistic, aggregated using the maximum sum of significant adjacent samples, *p*_*sample*_<.05), and followed up any significant effects by collapsing across significant timepoints and comparing individual conditions with paired-samples *t*-tests and Bayes Factors (uniform prior, lower bound: −0.3 a.u., upper bound: 0.3 a.u.).

### Univariate Electrode Sensitivity

To determine which electrodes were most informative for the forward encoding analyses, we tested the sensitivity of each electrode to both orientation and mismatch information (*Fig 3C and 3F*). The baseline-corrected signal at each electrode and time point in the epoch was regressed against a design matrix that consisted of the sine and cosine of the variable of interest (orientation or mismatch), and a constant regressor [30]. We calculated sensitivity, *S*, using the square of the sine (β_SIN_) and cosine (β_COS_) regression coefficients:

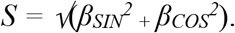

*S* was normalised against a null distribution of the values expected by chance. The null distribution was computed by shuffling the design matrix and repeating the analysis 1000 times. The observed (unpermuted) sensitivity index was ranked within the null distribution (to produce a p-value) and z-normalised using the inverse of the cumulative Gaussian distribution (*μ* = 0, *σ* = 1). The topographies shown in *Fig 3C and 3F* reflect the group averaged z-scores, averaged across each time period of interest.

---

**S1 Fig.**
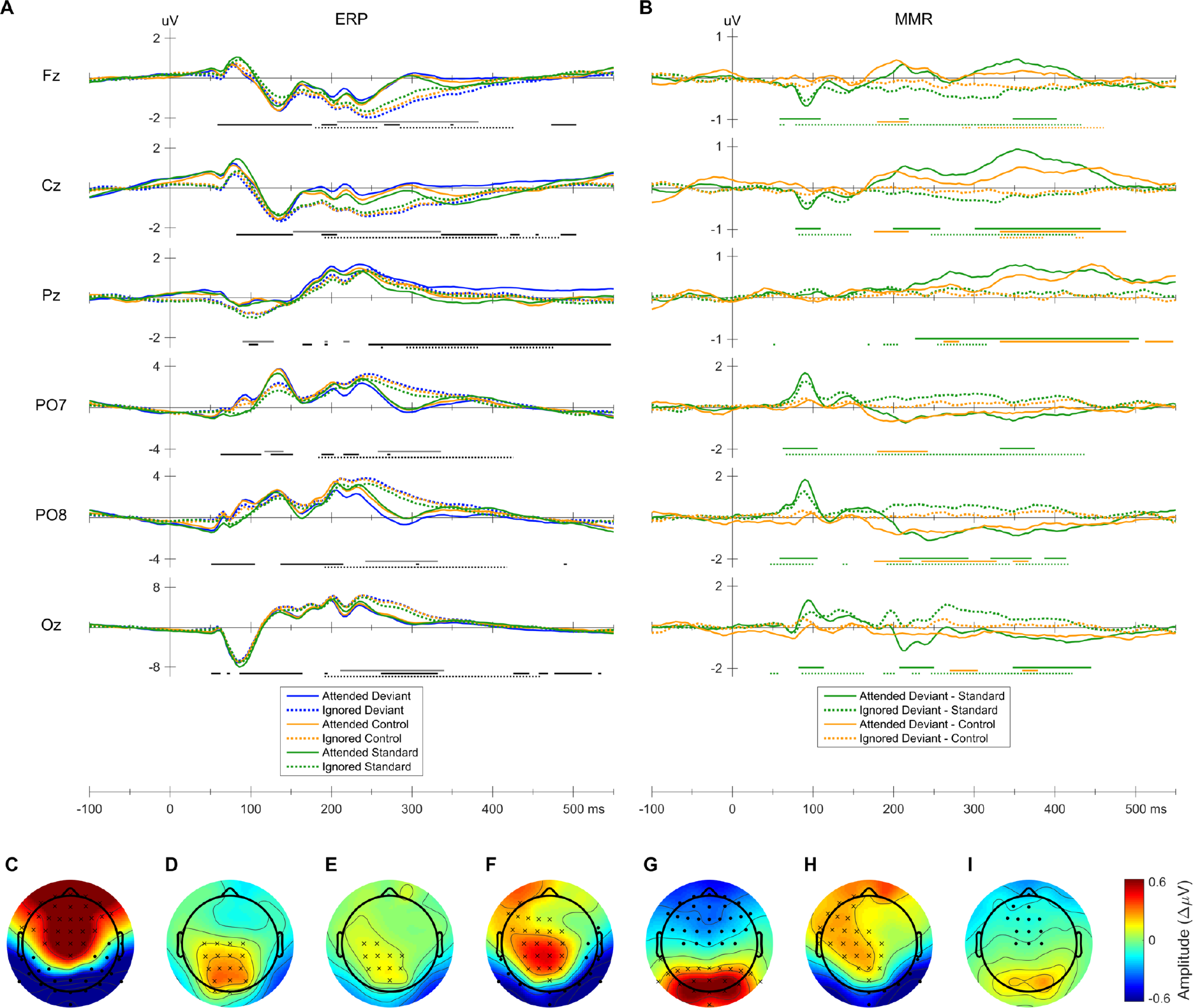
Event-related potentials (ERPs) and mismatch responses (MMRs). **(A)** ERPs at selected electrodes, shown separately for each condition. Bars underneath each plot indicate time points at which there was a significant main effect of attention (solid grey bar), significant main effect of prediction (solid black bar), or a significant interaction between attention and prediction (dotted black bar) at the plotted electrode. **(B)** Classic MMR (deviants - standards) and genuine MMR (deviants – controls) at selected electrodes, plotted separately for each level of attention. Green and yellow lines denote the classic MMR and genuine MMR, respectively; solid and dashed lines denote attended and ignored stimuli, respectively. Bars underneath each plot indicate timepoints at which there was a significant MMR in the corresponding condition, at the plotted electrode. Attended deviants were significantly different from attended standards (39 – 504 ms, cluster-corrected *p* < .001) and attended controls (172 – 550 ms, cluster-corrected *p* < .001). Ignored deviants were significantly different from ignored standards (47 – 438 ms, cluster-corrected *p* < .001) and ignored controls (285 – 461 ms, cluster-corrected *p* = .001) **(C-I)** Topographies of effects collapsed across time points between 200 and 300 ms. Asterisks and dots denote electrodes with larger, or smaller responses, respectively, in at least 25% of the displayed time points. **(C)** Main effect of attention (attended – ignored). **(D)** Classic MMR (deviants – standards). **(E)** Genuine MMR (deviants - controls). **(F)** Classic MMR during the grating task (attended deviants – attended standards). **(G)** Classic MMR during the dot task (ignored deviants – ignored standards). **(H)** Genuine MMR during the grating task (attended deviants – attended controls). **(G)** Genuine MMR during the dot task (ignored deviants – ignored standards).

**S2 Fig.**
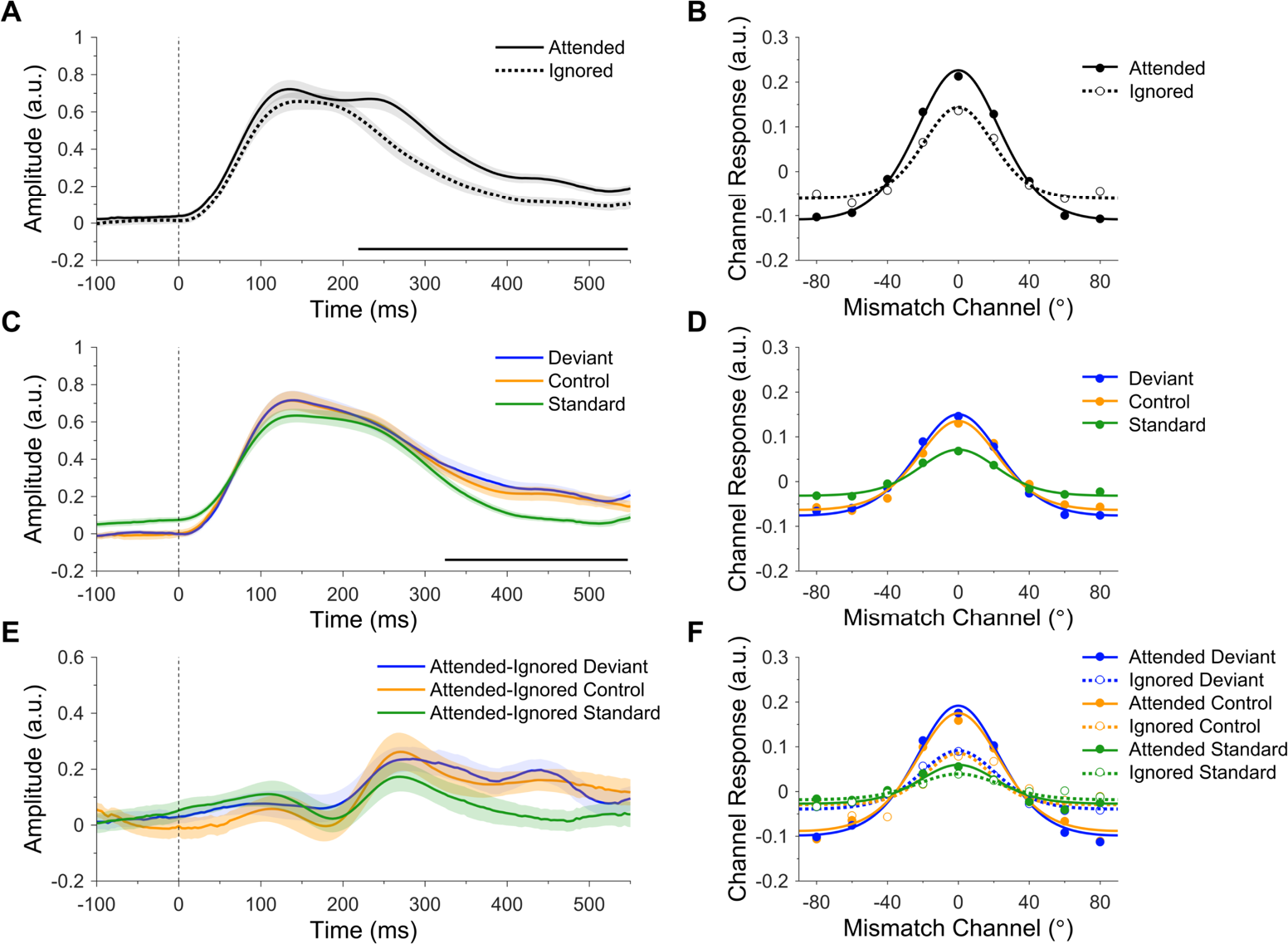
Independent main effects of attention and prediction on orientation response profiles, showing standards, deviants, and controls. **(A)** Main effect of attention on orientation response profiles. The amplitude of attended gratings was larger than that of ignored gratings (219 - 550 ms, cluster-corrected *p* = .001). Shading denotes standard error of the mean. The black bar along the x-axis denotes significant time points. **(B)** Orientation response profiles, collapsed across significant time points in **A**. Dots show activation in each of the nine modelled orientation channels. Curved lines show the functions used to quantify the amplitude and concentration of orientation-tuned responses (fitted to grand average data for illustrative purposes). **(C)** Main effect of prediction on orientation response profiles (black bar along the x-axis denotes significant time points, 324 – 550 ms, cluster-corrected *p* < .001). The amplitude of standards was reduced relative to both deviants and controls. **(D)** Orientation response profiles, collapsed across significant time points in **C**. **(E)** Interaction between attention and prediction on orientation response profile amplitude. Time-courses show the effect of attention (attended – ignored) on each stimulus type. **(F)** Orientation response profiles, collapsed across time points in the non-significant but trending cluster in **E** (414 - 481 ms, not displayed, cluster-corrected *p* = .093).

**S3 Fig.**
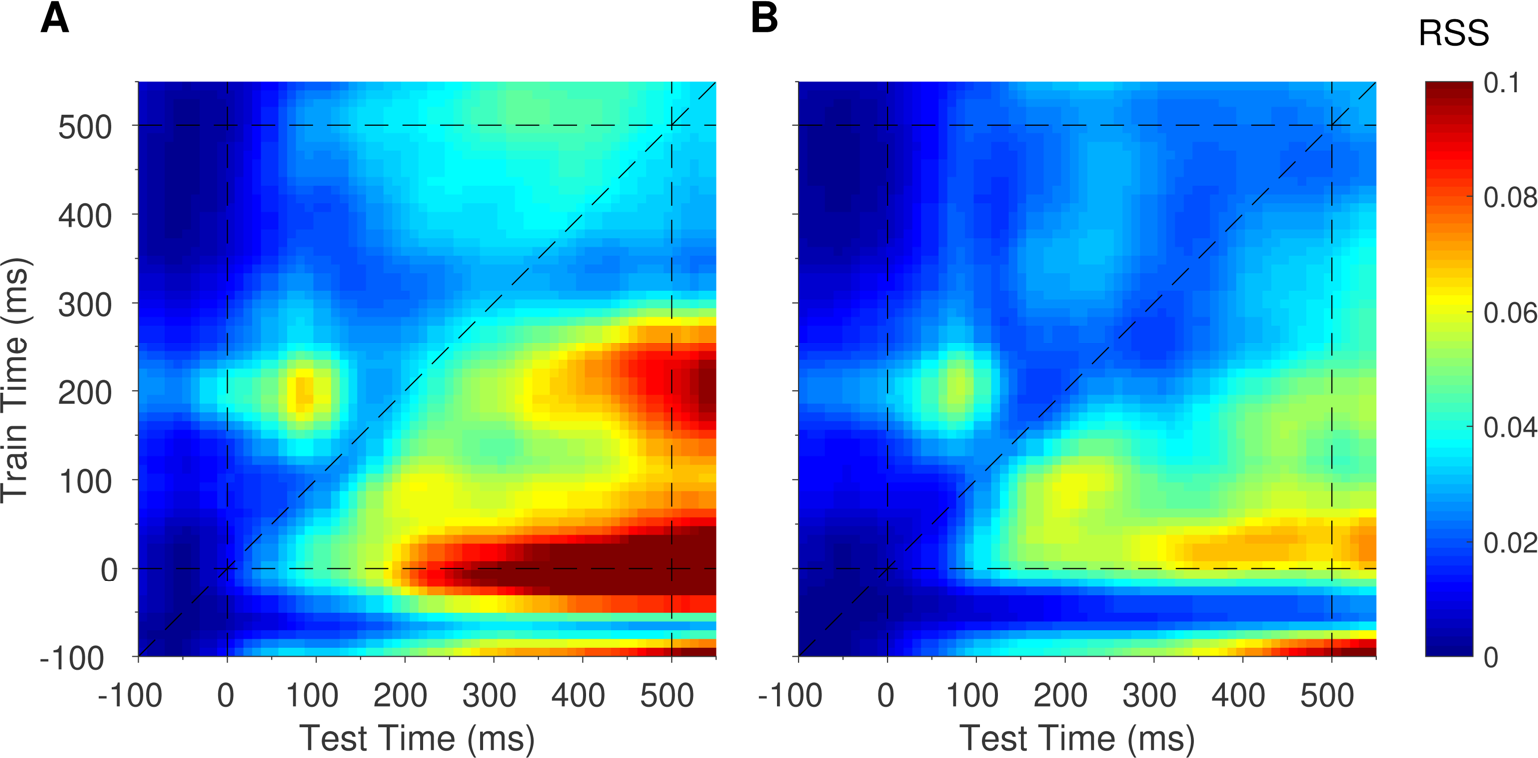
Residual sum of squares (RSS) for exponentiated cosine functions fitted to generalised mismatch response profiles (*Fig 5*). Note the high RSS values along the x-axis beginning at 200 ms, indicating that the apparent generalisation of spatial maps trained at stimulus onset to later times in the epoch (*Fig 5*, red patch along the x-axis) was likely due to noise.

## Notes

This research was supported by the Australian Research Council (ARC) Centre of Excellence for Integrative Brain Function (ARC Centre Grant CE140100007). J.B.M was supported by an ARC Australian Laureate Fellowship (FL110100103). M.I.G. was supported by a University of Queensland Fellowship (2016000071). The authors declare no competing financial interests.

